# Risky bait: capture experience alters feeding mode toward angling gear in red sea bream

**DOI:** 10.64898/2026.09.10.750600

**Authors:** Kohji Takahashi

## Abstract

When food resources provide reward while simultaneously imposing risk, animals can regulate their behaviour to continue exploiting them while reducing associated costs. Angling presents such a feeding situation, in which obtaining bait can result in hooking and capture. Although fish become less vulnerable to angling following capture experience, the behavioural changes underlying this reduced vulnerability remain poorly understood. Here, we used fine-scale video observations to examine feeding behaviour of red sea bream (*Pagrus major*) toward angling gear before and after capture experience. Naïve fish rapidly swallowed the bait and were subsequently captured, whereas capture-experienced fish continued to approach and feed on the gear but shifted from swallowing to biting. This feeding-mode shift may allow fish to maintain exploitation of the bait while reducing capture risk. Our observations provide a behavioural explanation for reduced angling vulnerability and suggest that direct experience with capture risk can alter how fish exploit a rewarding food resource.

## Introduction

Foraging behaviour in animals is determined by the trade-off between resource acquisition and associated risk [1,2]. In some ecological situations, however, a resource can carry value as both reward and risk, e.g., toxic prey or prey associated with injury risk can serve as both food and a source of risk. When food simultaneously carries both reward and risk, avoiding it results in the loss of feeding opportunities, whereas exploiting it exposes animals to risk [3].

When encountering food resources associated with risk, animals can regulate their exploitation by modifying resource choice, foraging intensity, and fine-scale feeding behaviour [4-6]. For example, birds experienced with toxic prey regulate their consumption of such prey according to their current toxin burden [7]. Such behavioural flexibility can allow animals to continue exploiting these resource while reducing their associated risks.

Angling provides a distinctive case of a risky food resource for fish because the capture risk is generated by angling gear rather than being an intrinsic property of the food. For naïve fish, baited angling gear initially provides a food reward, whereas capture experience provides information about the associated risks of hooking and capture. Thus, unlike responses to intrinsically dangerous food, behavioural responses to angling gear can be modified through direct experience with the associated anthropogenic risk. However, how fish reorganize their behaviour toward such a resource after experiencing its risk remains unclear.

Previous studies have shown that fish become less vulnerable to angling following capture experience, consistent with avoidance learning toward angling gear [8−10]. However, these studies have almost exclusively quantified capture success rather than the behavioural responses underlying reduced vulnerability.

We investigated angling gear avoidance learning in individually tested red sea bream *Pagrus major* juveniles [11]. This species rapidly reduces vulnerability to angling after only one or two angling experiences. Surprisingly, fish continued feeding on the bait throughout repeated presentations, with pecking behaviour even increasing despite their reduced vulnerability to angling. This observation raised the possibility that reduced angling vulnerability might involve changes in how fish exploit the bait after experiencing its associated risk, rather than simply avoidance of the gear.

In this study, we therefore used fine-scale video observations to examine how capture experience alters feeding behaviour toward angling gear in red sea bream. By comparing behaviour before and after capture, we examined how fish regulate their interaction with a rewarding resource associated with human-generated capture risk and aimed to identify the behavioural process underlying reduced angling vulnerability.

## Methods

### Fish

To examine fine-scale feeding behaviour toward angling gear, individual *P. major* were placed singly in small experimental tanks for observation of their feeding behaviour. Fish used in the experiment were hatchery-reared juveniles obtained from a commercial aquaculture strain and raised in a closed recirculating system in a rearing tank prior to the experiment. Four individuals were used in the experiment, and they were naïve to angling gear prior to the experiment (standard length: 62‒70 mm).

### Experimental setup

Experimental observations were conducted in tanks (length 27 × width 19 × height 15 cm). Each tank was covered with blue plastic sheets on all sides except for the observation side, and filled with 7 L of seawater identical to that used in the rearing tank. Aeration and a heater were installed to maintain the water temperature at approximately 24°C. Tanks were illuminated from the front. A video camera was positioned in front of the uncovered side of the tank to record fish behaviour. Four tanks were arranged side by side, and the entire experimental setup was covered with a sheet to minimize disturbance from the experimenter.

### Experimental procedure

Fish were randomly netted from the rearing tank and individually introduced into the experimental tanks, where they were acclimated overnight. On the following day, pellets were provided, and angling trials were initiated immediately after feeding was confirmed.

The angling gear was set up in the same manner as in a previous study [11](Sup Fig. S1). Briefly, the gear consisted of a rod, a fishing leader tied at one end to the tip of the rod, a fishing sinker attached to the other end, and a single hook (No. 2.5, SASAME HOOKS. LTD., approximately 8 mm in height) connected to a 5 cm length of 0.13mm nylon line. In this experiment, crushed hermit crab was used as bait.

At the beginning of each trial, approximately 5 pellets were presented to confirm feeding motivation. Subsequently, the angling gear was presented from behind the cover. When the fish appeared to take the bait in its mouth, the rod was lifted in an attempt to hook the fish. If the fish remained hooked to the line, it was immediately lifted above the water surface. If the fish dropped during lifting, the bait and gear were adjusted and the gear was reintroduced into the tank, after which behavioural observation resumed. For captured fish, the hook was removed above the water, and the fish was returned to the tank.

The angling gear was presented again 60 min after capture. This sequence constituted one trial, and 3‒4 trials were conducted for each individual. The experiment was recorded with a video camera, and behavioural changes toward the angling gear before and after capture experience were analyzed from the recordings. The data from Trial 2 of Individual 1 were excluded due to a recording failure. For Individual 4, the experiment was terminated because no immediate feeding response to pellets was observed from the second presentation of the angling gear.

### Behavioural analysis

From the recorded videos, we measured feeding latency, feeding behaviour, distance to the angling gear, and capture success for each individual and trial. The latency was defined as the time from presentation of either pellets or angling gear to the first feeding response. Feeding behaviour toward the angling gear was operationally classified into two categories: “swallowing”, in which the bait and hook were fully taken into the mouth, and “biting”, in which the fish pecked at the bait without taking the hook into the mouth. Additionally, when a fish was not captured during lifting, the event was classified as either a “drop”, in which the hook came off after the fish surfaced, or a “spit-out”, in which the fish spit out the hook underwater. We recorded the time at which each behaviour occurred and the total number of occurrences within a 60 s period. Distance to the angling gear was measured by calculating the distance from the bait on the gear to the tip of the snout on the video screen every second, and the distance was calibrated using an on-screen scale. Distance measurements were taken from the presentation of the gear until the fish was caught or until 60 s had elapsed. Each trial was categorized based on whether the fish was captured during the 60 s presentation period, and the behavioural patterns described above were compared between capture and non-capture trials. Given the small number of individuals, behavioural patterns were described exploratorily rather than subjected to formal statistical inference.

## Results

Before capture experience, all individuals were captured in the initial trial. Feeding latency to pellets and the angling gear was 1 ± 0 s and 2‒5 s, respectively, indicating rapid responses to both stimuli (Sup Table S1). The number of feeding events toward the angling gear ranged from 1 to 4, and all feeding events consisted of “swallowing” behaviour. Individuals No.1 and 3 were captured immediately after the first swallowing, resulting in only a single feeding event (Fig. 1a). Individual No. 2 showed multiple instances of “drop”; however, all drops occurred after the fish had been lifted above the water surface, and all subsequent feeding also consisted of swallowing. Distance to the angling gear rapidly decreased immediately after gear presentation (Fig. 2a, Sup Fig. S2, Sup Video 1); i.e., the fish approached the gear, fed immediately, and were subsequently captured.

**Fig. 1.**
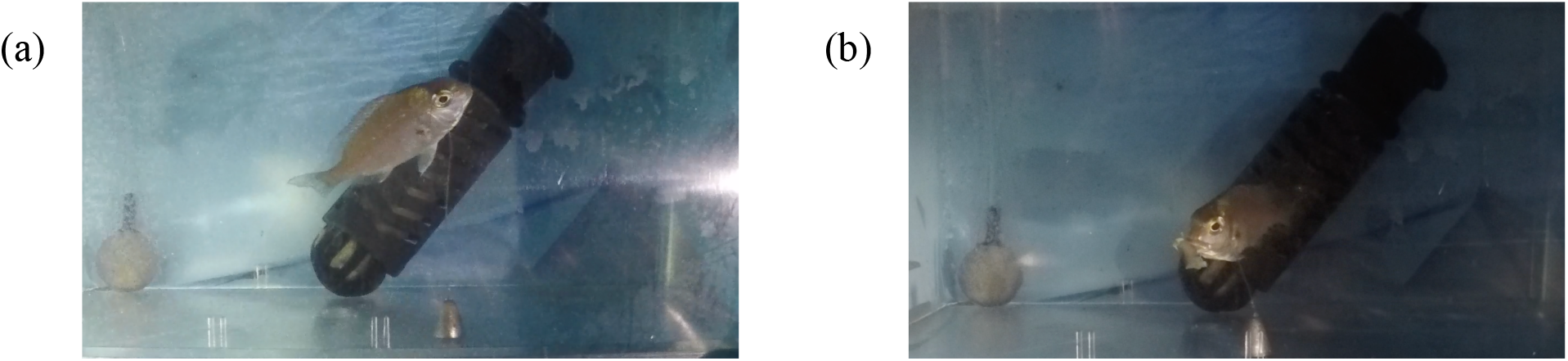
Feeding behaviour toward angling gear in red seabream. Representative video frames showing “swallowing” in Trial 1 (a) and “biting” in Trial 3 (b) from Individual No. 1.

**Fig. 2.**
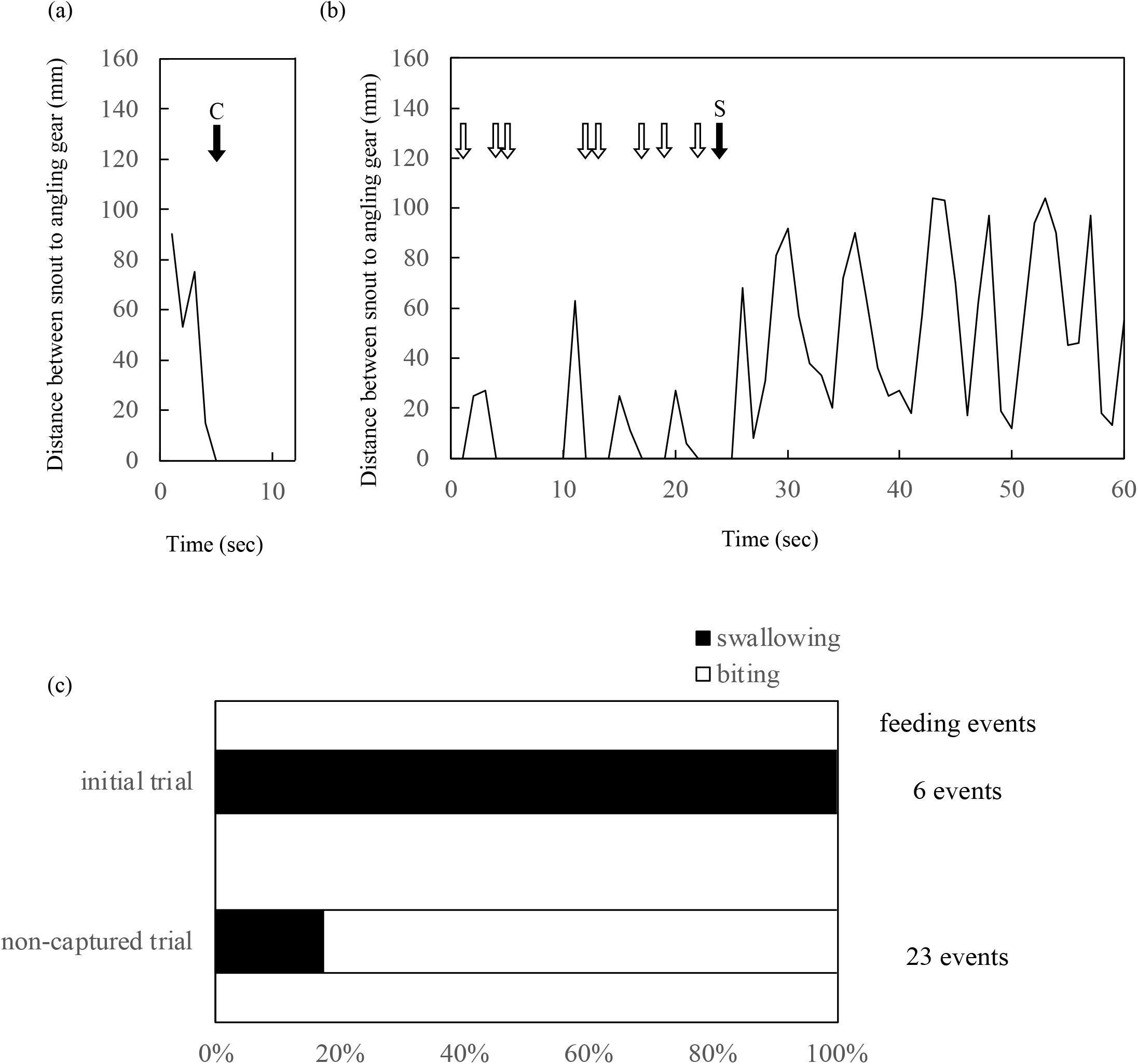
Temporal changes in distance between the fish snout and angling gear in individual No. 1. (a) Initial trial before capture experience and (b) non-capture trial following capture experience. White arrows indicate biting and black arrows indicate swallowing, C, captured; D, Drop; S, Spit out. (c) Proportions of swallowing and biting events in initial and non-capture trials pooled across all individuals. Number on the right indicate the total number of feeding events.

Following one or more capture experiences, all individuals eventually completed a trial without being captured during the 60 s presentation period. Even in non-captured trials, feeding occurred within a few seconds, with latencies of 0‒2 s for pellets and 2‒5 s for the angling gear (Sup Table S1). Only in Trial 3 of No. 2 did the individual immediately feed the pellets but show no feeding on the gear within the 60 s period. However, the same individual again fed on the gear immediately after presentation in Trial 4.

In non-captured trials, the numbers of feeding events ranged from 5 to 10, and approximately 90% of these consisted of “biting” behaviour (Fig. 1b, Sup Video 2). Even when swallowing occurred, fish exhibited “spit-out” behaviour in all cases, resulting in no capture within the 60 s period.

In non-captured trials, distance to the angling gear fluctuated dynamically over time (Fig. 2b, Sup Fig. S2). Fish repeatedly approached and withdrew from the gear, exhibiting biting during these movements. Additionally, in Trial 3 of No. 2, the fish did not feed but maintained a constant distance while orienting toward the gear. However, the same individual exhibited repeated approach-withdrawal movements in Trial 4. Although the trial at which behavioural changes first appeared differed among individuals, all individuals eventually exhibited similar behavioural patterns during non-capture trials.

## Discussion

Capture experience altered the feeding mode of red sea bream toward angling gear. Naïve fish rapidly swallowed the bait immediately after gear presentation and were subsequently captured. In contrast, fish with capture experience continued to approach and feed readily on the gear, but shifted from swallowing to biting. Even when swallowing occurred, fish frequently spat out the bait before capture. These observations suggest that capture experience did not substantially suppress feeding motivation. Instead, it altered how fish fed rather than whether they fed, shifting their feeding mode from swallowing to biting.

Behavioural changes were also evident in movement patterns toward the angling gear. Following capture experience, fish eventually exhibited repeated approach‒withdrawal movements from the gear, with the distance to the gear fluctuating dynamically over time. Biting frequently occurred during these movements. Thus, behavioural changes were not restricted to feeding mode: fish continued to interact with the gear rather than simply avoiding it.

Previous studies showed that angling vulnerability can be reduced through capture experience [8-11]. However, the present observations suggest that reduced angling vulnerability cannot be explained simply by avoidance of the gear. Instead, fish continued to interact with the gear while modifying both their movement patterns and feeding mode. In particular, shifting their feeding mode from swallowing to biting may allow fish to continue exploiting the bait while reducing the likelihood of capture. Reduced angling vulnerability may reflect changes in how fish exploit angling gear. By examining fine-scale behaviour toward angling gear, the present study provides a behavioural explanation for reduced angling vulnerability that could not be inferred from capture success alone.

The behavioural changes observed after capture experience are consistent with behavioural regulation toward a risky food resource [4-7]. Risky prey exploitation may involve adjustments at multiple stages of interaction, including approach, feeding, and rejection [3]. In the present study, capture experience was followed by repeated approach‒withdrawal movements, a shift from swallowing to biting, and rejection of swallowed bait. Together, these changes suggest that fish regulated multiple aspects of their interaction with angling gear in a human-generated risky feeding situation that they can repeatedly encounter.

Among the behavioural regulations observed in the present study, the shift in feeding mode from swallowing to biting was particularly notable. Although experience-dependent changes in sampling behaviour toward risky food have been reported in fish [12], the present shift from swallowing to biting suggests that capture experience can also alter the feeding decision of how to exploit the food. Such a shift may be particularly functional in an angling context because swallowing the baited hook directly increases the likelihood of capture. The present study suggests that behavioural regulation toward risky food can be shaped by direct experience with risk and can involve changes in how the resource is exploited while interaction with the resource is maintained.

The behavioural changes were clear and consistently observed in all individuals. Nevertheless, this study was intended as an exploratory behavioural description based on a limited number of individuals. It will be necessary to verify the generality of the behavioural patterns described here through larger-scale quantitative analysis. Future studies examining repeated approach-withdraw behaviour together with fine-scale behavioural variables, such as gaze direction, body posture, and movement dynamics, will help clarify the behavioural mechanisms underlying decision-making toward stimuli carrying both reward and risk.

## Notes

### Competing Interest Statement

The authors have declared no competing interest.

