## Supplementary material for "Risky bait: capture experience alters feeding mode toward angling gear in red sea bream": Sup figs1, figs2, tables1

Supplemental File

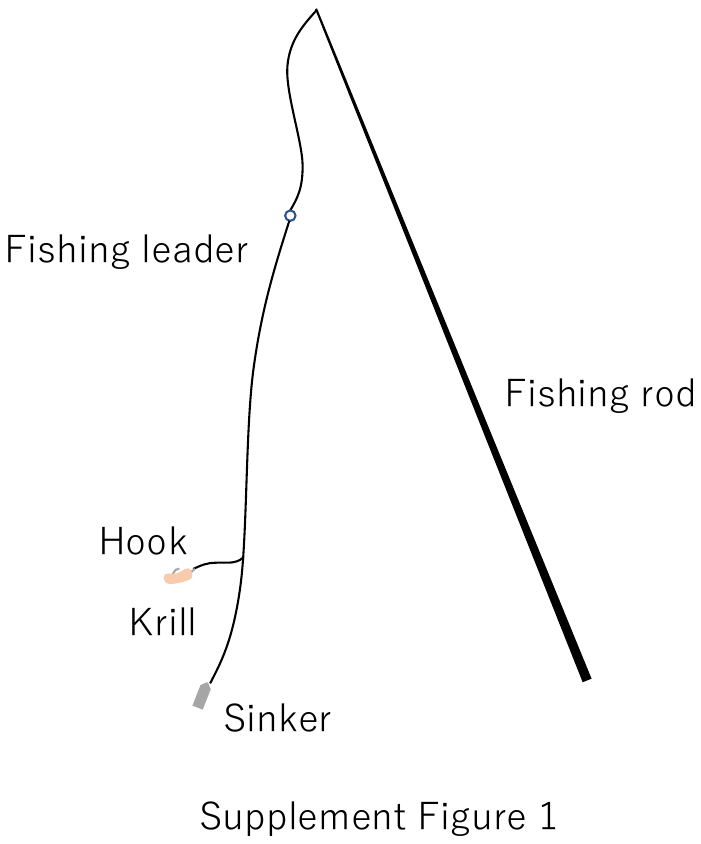

Figure S1 Schematic illustration of angling gear

Table S1 Datum of individuals in Experiment

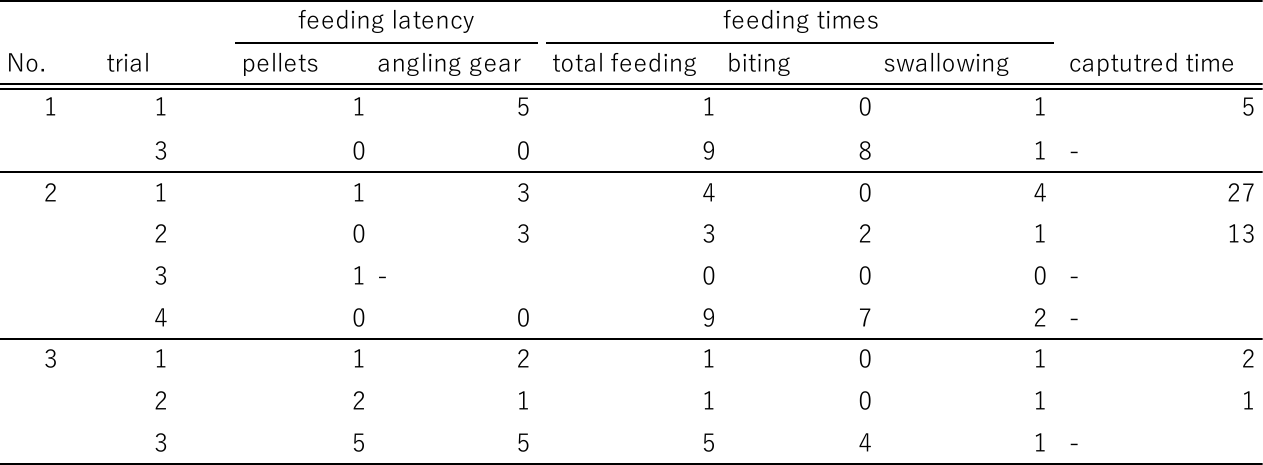

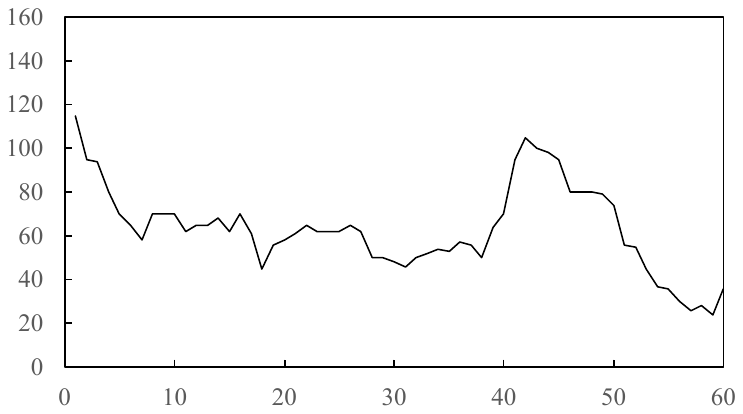

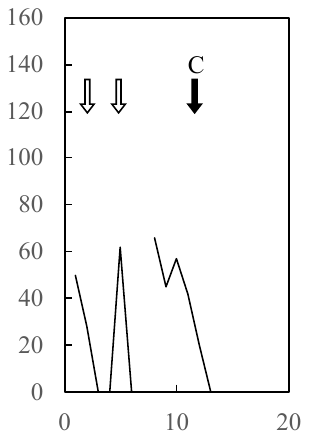

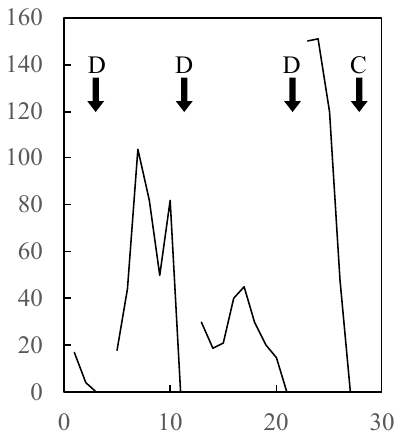

Distance between snout to angling gear (mm)

Trial 4

Trial 3

Trial 2

Trial 1

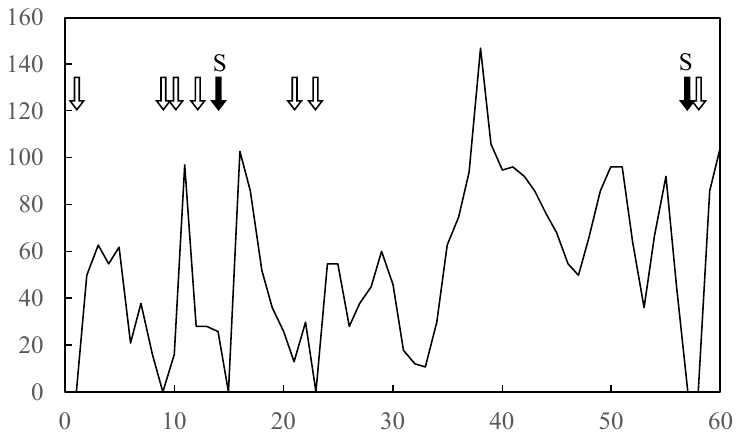

Time (sec)

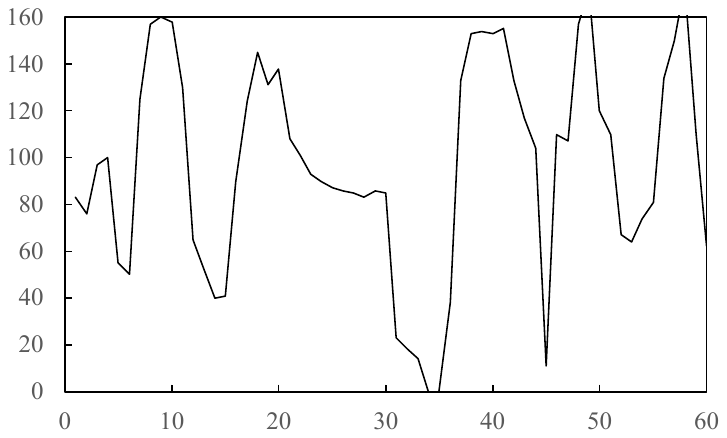

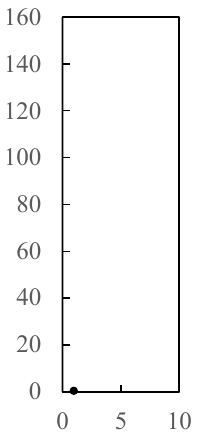

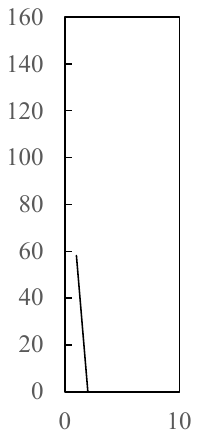

Trial 2

Trial 1

Trial 3

Figure S2 Temporal changes in distance between the fish snout and angling gear in individual No. 2 (upper) and No.3 (lower). White arrows indicate biting and black arrows indicate swallowing, C, captured; D, Drop; S, Spit out.

Time (sec)
